# *Drosulfakinin* signaling encodes early-life memory for adaptive social plasticity

**DOI:** 10.1101/2024.04.08.588562

**Authors:** Jiwon Jeong, Kujin Kwon, Terezia Klaudia Geisseova, Jongbin Lee, Taejoon Kwon, Chunghun Lim

**Author notes:** These authors contributed equally (Jiwon Jeong, Kujin Kwon). Correspondence (T.K.), (C.L.).

## Abstract

*Drosophila* establishes social clusters in groups, yet the underlying principles remain poorly understood. Here we performed a systemic analysis of social network behavior (SNB) that quantifies individual social distance (SD) in a group over time. The SNB assessment in 175 inbred strains from the Drosophila Genetics Reference Panel revealed a tight association of short SD with long developmental time, low food intake, and hypoactivity. The developmental inferiority in short-SD individuals was compensated by their group culturing. By contrast, developmental isolation silenced the beneficial effects of social interactions in adults and blunted the plasticity of SNB under physiological challenges. Transcriptome analyses showed genetic diversity for SD traits, whereas social isolation reprogrammed select genetic pathways, regardless of SD phenotypes. In particular, social deprivation suppressed the expression of the neuropeptide Drosulfakinin (*Dsk*) in three pairs of adult brain neurons. Male-specific DSK signaling to Cholecystokinin-like receptor 17D1 mediated the SNB plasticity. In fact, transgenic manipulations of the DSK signaling were sufficient to imitate the state of social experience. Given the functional conservation of mammalian *Dsk* homologs, we propose that animals have evolved a dedicated neural mechanism to encode early-life experience and transform group properties adaptively.

## Introduction

Animals interact with other individuals in distinct social environments (1-3). For instance, a pair of animals display aggression or mating behaviors, whereas a group of individuals may show collective behaviors likely through a network of social interactions. Social experience further impacts the physiological properties of individuals, such as feeding, sleep/circadian rhythms, aggression, stress response, and longevity (4-10). Consequently, social network behavior (SNB) is thought to confer group fitness and thus is conserved across species (3, 11). Nonetheless, it remains elusive how group properties have evolved with other individual traits and how animals process social memory to shape their physiology. *Drosophila* has long been considered a solitary species. Yet a group of flies can display a distinct property of SNB under defined experimental conditions (12-17), serving as an ideal genetic model to address our questions above. A list of biophysical parameters has been measured to quantify group or individual properties, collectively defining social interaction criteria in *Drosophila* (2, 18). Although social network measurements comprehensively describe group behaviors, we focused on the clustering property in a group of flies (13, 19). Social clustering is an intuitive measure that integrates diverse social interactions via multiple sensory cues (12, 18, 19), accompanying reductions in both social distance (SD) among group members and their moving speed over time. Simple SD assessment facilitated our approaches that aligned large-scale datasets for group behaviors to physiological traits, differential gene expression, and neurogenetic manipulations, thereby elucidating the principles of SNB and its plasticity.

## Results

### SNB is a quantitative trait in a natural *Drosophila* population

We employed the *Drosophila* Genetic Reference Panel (DGRP) to determine whether SNB has evolved with specific genetic factors and physiology. The DGRP consists of approximately 200 inbred wild-type strains and functions as a practical genetic library to explore the correlation between naturally occurring genetic variations and complex animal behaviors (20-22). We video-recorded a group of 16 male flies freely moving in an open arena for 10 min and quantitatively assessed their group properties over time (Fig. 1A). These included SD between individual group members (Fig. 1B), walking speed, and the centroid velocity of a given group. The DGRP lines displayed a range of distributions for the three parameters (Fig. 1C), and we found their significant correlations among 175 DGRP lines (Fig. 1D). For instance, short-SD lines exhibited slow walking speeds and low centroid velocities (e.g., DGRP73, DGRP 563, and DGRP370), which eventually led to the formation of a stable cluster within the arena (Fig. 1D and S1). By contrast, long-SD lines had fast walking speeds and high centroid velocities (e.g., DGRP360, DGRP707, and DGRP317). The locomotion trajectories of individual flies further confirmed these characteristics (Fig. 1B and 1E). Short-SD flies gradually reduced SD over time and stayed in the cluster, whereas long-SD flies persistently explored the arena and sustained “social distancing”. These results provide convincing evidence that SD is an inheritable group trait from natural *Drosophila* variants. We subsequently selected the top three DGRP lines representing either short- or long-SD phenotypes to elucidate the physiological significance of *Drosophila* SNB and its underlying mechanisms.

**Fig. 1.**
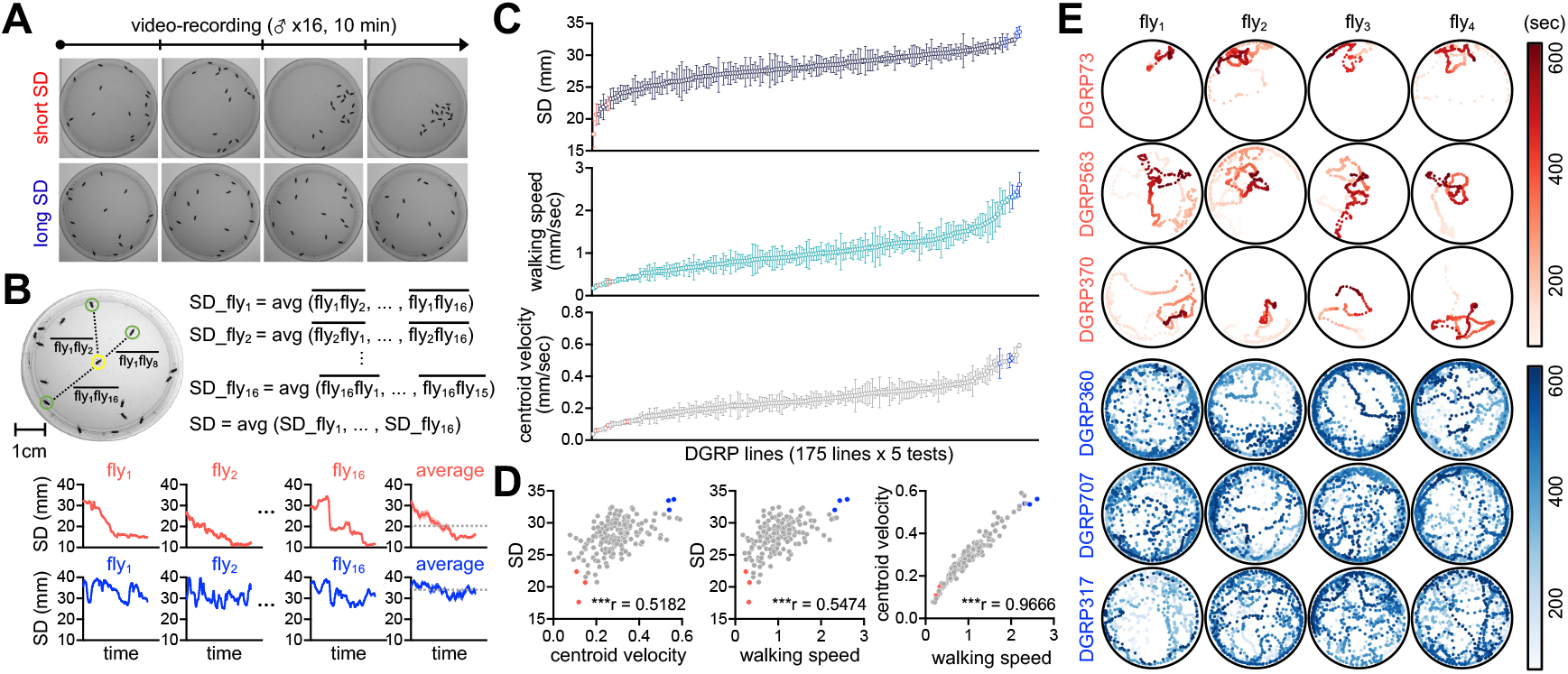
Social network behavior is a quantitative trait in a natural *Drosophila* population. (A) 10-min video recording of social network behavior (SNB) in a group of 16 male flies. (B) The definition of social distance (SD). (C) Quantitative assessment of SNB by ranking SD, walking speed, and centroid velocity among 175 DGRP lines. Data represent means +/- SEM (*n* = 5). (D) Significant correlation among SD, walking speed, and centroid velocity. ^***^*P* < 0.001, as determined by Spearman correlation analysis. (E) 10-min locomotion trajectories of individual flies from the three representative DGRP lines displaying short (red) or long SD (blue).

### Social interactions compensate for developmental inferiority in short-SD larvae

Why do flies display SNB? One clue comes from the previous observation that *Drosophila* larvae collectively dig culture media and improve food accessibility, possibly facilitating their constitutive feeding during early development (15). In fact, we found that the short-SD lines had higher numbers of larvae per cluster than the long-SD lines (Fig. 2A). These observations suggest that *Drosophila* express SNB traits from early development, and the sociality persists in adults. To better define the social-interaction effects on *Drosophila* physiology, we obtained socially enriched or deprived larvae from fertilized eggs (Fig. 2B) and then compared their developmental phenotypes among DGRP lines. Socially isolated larvae from the short-SD lines displayed poorer digging activity and lower food intake than those from the long-SD lines (Fig. 2C and 2D), which likely led to longer developmental time (Fig. 2E). Intriguingly, social isolation lowered the relative developmental success of the adult male progeny in the short-SD lines (Fig. 2F). Grouping of short-SD larvae improved their digging activity and food intake to levels observed in isolated long-SD larvae (Fig. 2C and 2D). The group culture also shortened developmental time and elevated male progeny ratio only in short-SD lines, making the properties of socially enriched groups comparable between short- and long-SD lines (Fig. 2E and 2F). Our group culture was not a competitive environment for limited resources to generally delay developmental time (23, 24). Yet these results indicate that the clustering property of short-SD lines may have evolved as a compensation mechanism for their developmental inferiority in individuals.

**Fig. 2.**
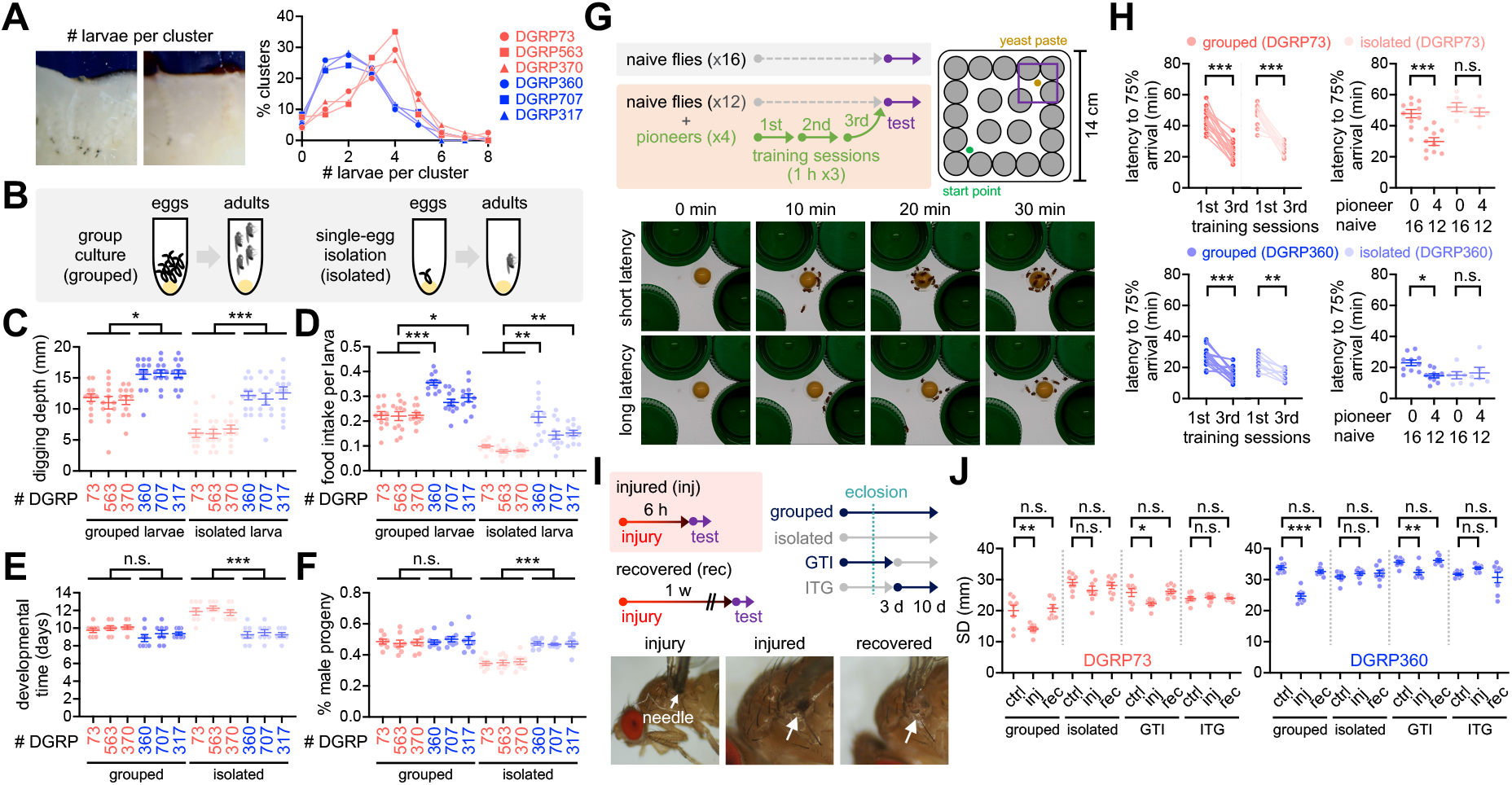
Early-life social experience confers beneficial effects on *Drosophila* development and physiology. (A) Larval clustering in short- (red) or long-SD lines (blue). % clusters were calculated from 30 vials. (B) Schematic for grouped vs. developmentally isolated culture conditions. (C and D) Grouped culture compensated for low food accessibility in the short-SD larvae. Data represent means +/- SEM (*n* = 12). ^*^*P* < 0.05, ^**^*P* < 0.01, ^***^*P* < 0.001, as determined by Kruskal-Wallis test with Dunn’s multiple comparisons test (digging depth, grouped), 1-way ANOVA with Tukey’s multiple comparisons test (digging depth, isolated; food intake, grouped), or Welch’s ANOVA with Dunnett’s T3 multiple comparisons test (food intake, isolated). (E and F) Grouped culture rescued developmental delay and low male-progeny ratio in the short-SD larvae. Data represent means +/- SEM (*n* = 8). n.s., not significant; ^***^*P* < 0.001, as determined by 1-way ANOVA with Tukey’s multiple comparisons test. (G) Experimental scheme for assessing social interactions in a maze assay. (H) Pioneer groups of short- (red, DGRP73) and long-SD lines (blue, DGRP360) were effectively trained in the maze assay, but social isolation blunted the pioneer effects on food-seeking behaviors in a group of naive flies. Data represent means ± SEM (*n* = 6□16). n.s., not significant; ^*^*P* < 0.05, ^**^*P* < 0.01, ^***^*P* < 0.001, as determined by 2-way repeated measures ANOVA with Sidak’s multiple comparisons test (training sessions) or 2-way ANOVA with Tukey’s multiple comparisons test (test sessions). (I) Experimental scheme for assessing injury-induced social plasticity. GTI, grouped-to-isolated culture transition; ITG, isolated-to-grouped culture transition. (J) Physical injury induced clustering behaviors in group-cultured but not developmentally isolated flies. Data represent means ± SEM (*n* = 8). n.s., not significant; ^*^*P* < 0.05, ^**^*P* < 0.01, as determined by 1-way ANOVA with Tukey’s multiple comparisons test.

### Early-life experience is necessary for social benefits on adult physiology and adaptive social plasticity

We further determined whether social interactions also benefit adult physiology in the short-SD lines. To this end, we designed a maze experiment where a group of flies were placed in a novel arena to determine how fast they could reach a food resource in the presence or absence of pre-trained colleagues (Fig. 2G). Our prediction was that social interactions between naïve and trained flies might facilitate their food-seeking, possibly mimicking social foraging in other species (25). We first confirmed that both short- and long-SD lines significantly shortened latency to the arrival of 75% of flies on food by iterative exposures to the maze (Fig. 2H and S2). A representative plasticity mutant of the *rutabaga* (*rut*) gene did not significantly shorten the arrival latency after three consecutive training sessions in our maze paradigm (Fig. S3A), implicating *rut*-dependent learning and memory in this process (26). The short-SD lines displayed poor performance in the maze assay as assessed by longer latency to the arrival of 75% naïve flies on food than the long-SD flies (Fig. 2H and S2). When combined with a group of naïve flies, the trained “pioneers” substantially improved the group property of food-seeking behaviors in both short- and long-SD lines (Fig. 2H and S2). The pioneer effects disappeared when the naïve group consisted of socially isolated individuals from egg development. These results support that social foraging in our maze paradigm is unlikely a simple collective response in adults, but it specifically requires early-life social experience.

The long-SD lines outcompeted the short-SD lines in both larval and adult assays, and the beneficial effects of their social experience were barely detectable. SNB in the long-SD adults was also insensitive to early-life experience, whereas the grouped culture promoted SNB in the short-SD lines (18) (Fig. S1B). We hypothesized that superior traits in long-SD individuals led to their genetic selection toward the degeneration of social interaction effects. Alternatively, long-SD genomes may still encode genetic programs for social activity, but they only express social traits under physiologically challenging conditions. Given the positive correlation between locomotion activity and SD among DGRP lines (Fig. 1D), we reasoned that injury could serve as a physiological cue to negatively impact locomotion and induce SNB even in the long-SD flies. We indeed discovered that mechanical injury shortened SD in both sociality types of DGRP lines (Fig. 2I, 2J, and S4A). The injury effects were transient because one week of recovery was sufficient to restore the original SD traits. Simple impairment of locomotor activities was unlikely to be responsible for the injury-induced SD reduction since the injury effects were not detectable in a group of socially isolated flies (Fig. 2J and S4A). We further found that an adult-specific grouping of developmentally isolated flies was insufficient to support the social plasticity. Finally, the *rut*-dependent memory pathway seems dispensable for developmental “social memory” since *rut* mutants displayed injury-induced social plasticity comparable to control flies (Fig. S3B). It remains yet to be determined whether or not the mechanical injury promotes SNB via a specific sensing mechanism. Nonetheless, these results suggest that individual strains differentially display social traits in adults; however, the potency of adaptive social plasticity is acquired through their early-life experience of social interactions.

### Unraveling the genetic basis of SNB and its plasticity

How is early-life social experience encoded in individual larvae to persist throughout development? To define the molecular signatures of social interaction phenotypes and their plasticity, we profiled differentially expressed genes (DEGs) in adult fly heads among distinct contexts of genetic and environmental sociality. The transcriptome analyses revealed significantly upregulated genes in the short- and long-SD lines (n genes = 191 and 199, respectively) as well as in grouped and socially isolated flies (n genes = 159 and 755, respectively) (Fig. 3A and 3B; Dataset S1 and S2). Gene ontology (GO) analyses showed no significant enrichment of specific GO terms in the short- or long-SD lines. We thus concluded that diverse genetic pathways led to social phenotypes in the given set of DGRP lines, as suggested previously (12). It is also consistent with the lack of the phylogenetic correlation between *Drosophila* species and their social network phenotypes (27). Nevertheless, select metabolic pathways were consistently upregulated in both types of DGRP lines by social isolation (7, 28) (Fig. 3C and 3D; Dataset S2 and S3).

**Fig. 3.**
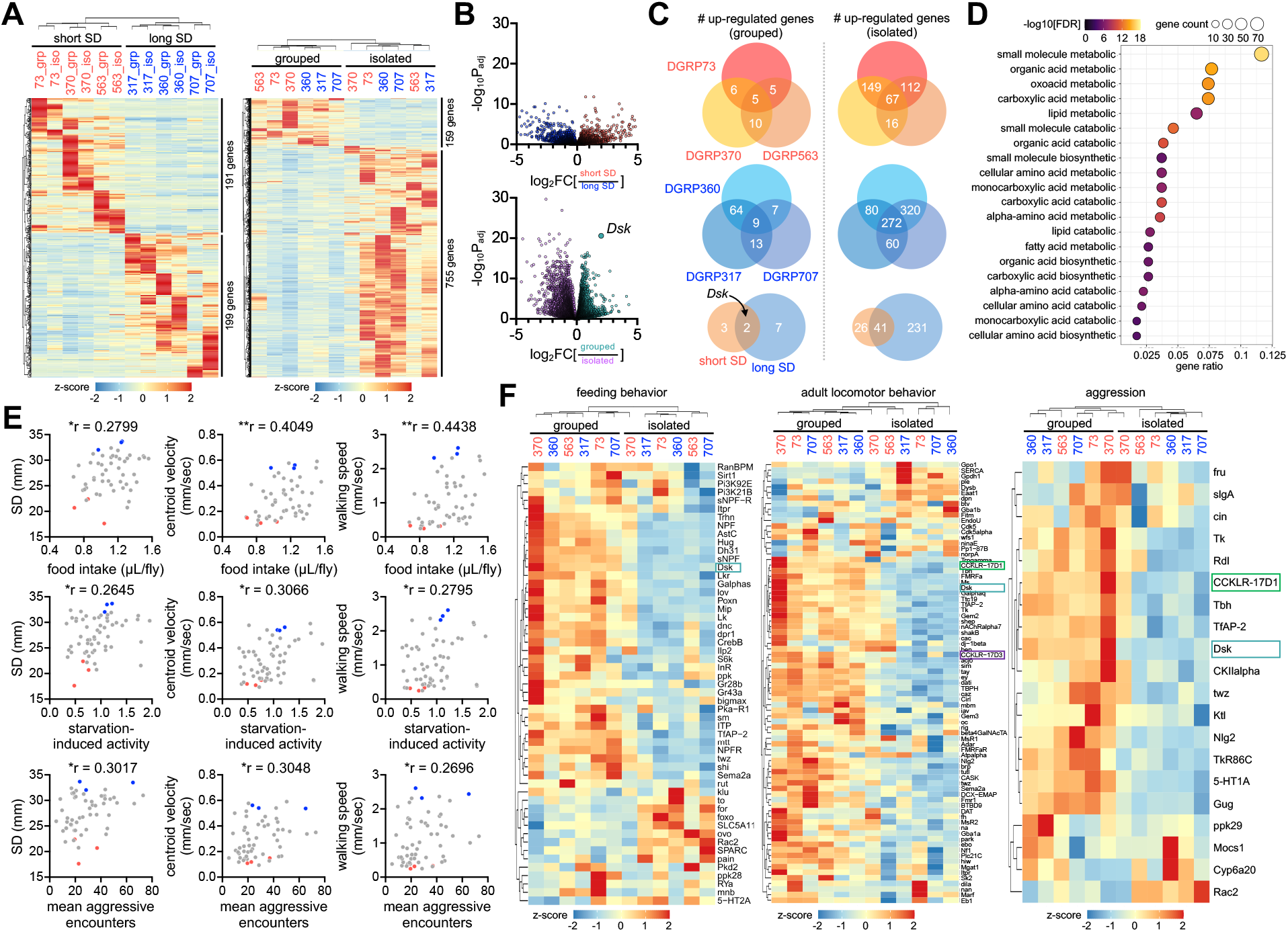
Social experience shapes gene expression profiles in *Drosophila* heads. (A) Heatmaps for differentially expressed genes (DEGs, > 2-fold difference with adjusted *P* < 0.05) in short-vs. long-SD lines (left); in grouped vs. isolated condition (right). Fly heads were harvested from grouped or isolated DGRP lines and their gene expression profiles were analyzed by RNA sequencing. Averaged counts per million were converted to z-score for visualization. (B) Volcano plots for DEGs in short-vs. long-SD lines (top); in grouped vs. isolated lines (bottom). Social interactions evidently upregulated the neuropeptide *Drosulfakinin* (*Dsk*) expression. (C) Overlapping DEGs in grouped vs. isolated condition across DGRP lines. *Dsk* was identified as a commonly upregulated gene by social interactions between short- (top, red) and long-SD lines (middle, blue). (D) Gene ontology analysis reveals upregulation of select metabolic pathways upon social isolation. (E) Significant phenotypic correlation of SNB to food intake, starvation-induced activity, and mean aggressive encounters among DGRP lines. ^*^*P* < 0.05, ^**^*P* < 0.01, as determined by Spearman correlation analysis. (F) Expression heatmap for genes implicated in feeding, adult locomotor behavior, and aggression. Downregulation of *Dsk* and its two receptors (CCKLR-17D1 and CCKLR-17D3) by social isolation was visualized in relevant gene categories.

Interestingly, the phenotypic alignment of DGRP lines revealed significant correlations of their social interaction behaviors to food intake, starvation-induced activity, and aggression, among others (29-31) (Fig. 3E). The association of multiple behaviors may indicate their overlapping evolution of regulatory genes and mechanisms. The expression heatmaps of relevant gene categories further visualized a subset of genes that displayed social experience-dependent expression across the DGRP lines analyzed (Fig. 3F). These indeed included *Drosulfakinin* (*Dsk*), a neuropeptide implicated commonly in aggression, food intake, satiety, and sexual behaviors (32-36). The mammalian *Dsk* homolog, cholecystokinin (CCK), plays a similar role in relevant physiology, indicating evolutionary conservation (37-39). In fact, *Dsk* and the two CCK-like receptors (i.e., *CCKLR-17D1* and *CCKLR-17D3*) were consistently downregulated upon social isolation (Fig. 3F). These observations prompted us to ask whether DSK signaling contributes to the plasticity of social behaviors through early-life experience.

### DSK neuron activity encodes early life experience for SNB plasticity

Immunostaining of whole-mount brains identified three groups of DSK-expressing neurons with distinct neuroanatomical morphology (i.e., MP1a, MP1b, and MP3) (32, 34) (Fig. 4A). The long-SD lines displayed relatively high DSK signals in the cell bodies compared to the short-SD lines (Fig. 4B). However, DSK levels in the neural projections were comparable between the two groups. Social isolation lowered DSK levels irrespective of the SD phenotype (Fig. 4A and 4B), consistent with our DEG analysis above, and the social-context effects were most evident in the MP1a neuron projections. Live-brain imaging of the genetically encoded Ca^2+^ indicator GCaMP showed that DSK neuron activity correlated with social experience and plasticity. Social deprivation reduced relative Ca^2+^ levels in DSK neurons, whereas mechanical injury generally elevated DSK neuron activity (Fig. 4C and 4D). Intriguingly, MP1a neuron activity did not respond to injury when transgenic flies were socially deprived during development, which may contribute to the lack of social behavior plasticity upon isolation. These observations are unlikely due to transgenic *Dsk*-Gal4 activity per se given that social isolation did not comparably affect the Gal4-dependent expression of a dendritic marker transgene in the DSK neurons (Fig. 4C and 4D).

**Fig. 4.**
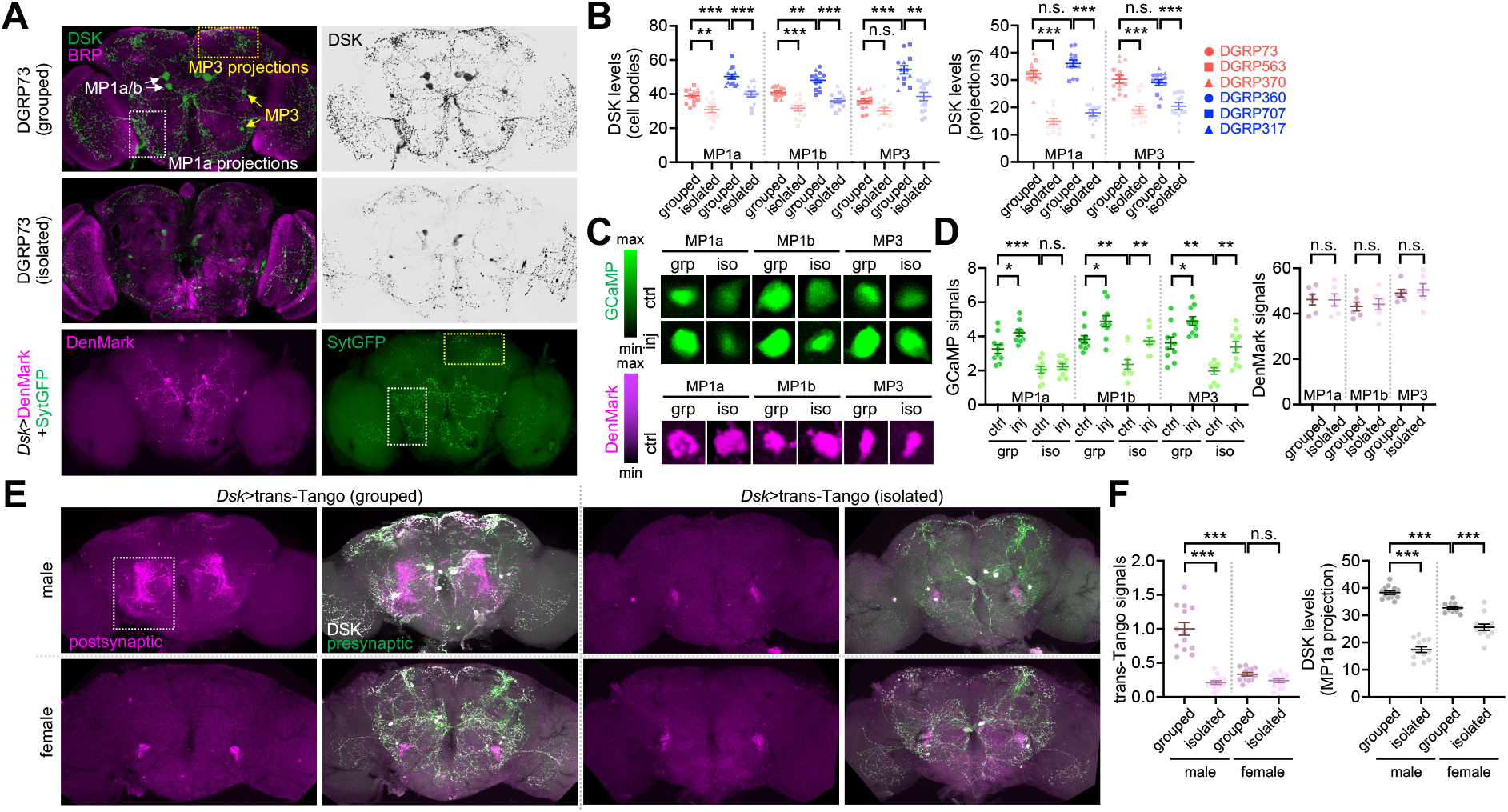
DSK neuron activity encodes social experience. (A and B) Social experience elevates DSK levels in the *Drosophila* brains. Whole-mount brains were co-immunostained with anti-DSK (green) and anti-BRUCHPILOT (BRP, a synaptic protein; magenta) antibodies. Dendrites (DenMark, magenta) and axons (SytGFP, green) of DSK neurons were independently visualized in a transgenic brain (*Dsk*>DenMark+SytGFP). The fluorescent DSK signals from confocal images were quantified using ImageJ. Data represent means ± SEM (*n* = 13). n.s., not significant; ^**^*P* < 0.01, ^***^*P* < 0.001, as determined by 2-way ANOVA with Tukey’s multiple comparisons test (MP1a/MP1b/MP3 cell bodies and MP1 projections) or aligned ranks transformation ANOVA with Wilcoxon rank sum test (MP3 projections). (C and D) Social experience and physical injury elevate DSK neuron activity. Social isolation masked an injury-induced Ca^2+^ increase in the MP1a neurons among other DSK neurons as assessed by the genetically encoded Ca^2+^ sensor GCaMP in live-brain imaging. The transgenic DenMark expression served as a negative control for social-isolation effects on GCaMP expression per se. Data represent means ± SEM (*n* = 10 for GCaMP; *n* = 6 for DenMark). n.s., not significant; ^*^*P* < 0.05, ^**^*P* < 0.01, ^***^*P* < 0.001, as determined by 2-way ANOVA with Tukey’s multiple comparisons test (GCaMP) or unpaired t-test (DenMark). (E and F) Social experience strengthens postsynaptic signaling of DSK neurons only in males. Postsynaptic partner of DSK neurons was visualized by the heterologous ligand-receptor signaling embedded in the transsynaptic mapping transgene (trans-Tango). The fluorescent trans-Tango signals from confocal images were quantified using ImageJ. Data represent means ± SEM (*n* = 13). Aligned ranks transformation ANOVA and 2-way ANOVA detected significant interactions of gender with trans-Tango signals and DSK levels in MP1a projections, respectively (*P* < 0.001). n.s., not significant; ^***^*P* < 0.001, as determined by Wilcoxon rank sum (trans-Tango signals) or Tukey’s multiple comparisons tests (DSK levels).

We also used the transsynaptic mapping transgene trans-Tango to visualize the postsynaptic partner of DSK neurons via heterologous ligand-receptor signaling (40, 41). Socially enriched, but not socially isolated, male flies displayed strong trans-Tango signals in the lateral protocerebrum where MP1a neuron projections are specifically enriched (32) (Fig. 4A, 4E, and 4F). DSK neurons have gender-specific presynaptic partners, which may mediate gender-specific signaling for sexual behaviors (32, 42). To our surprise, socially enriched female brains did not show trans-Tango signals as evidently as male brains (Fig. 4E and 4F). Moreover, social deprivation downregulated DSK levels in the MP1a projections less potently in females than in males (Fig. 4E and 4F). These observations suggest sexual dimorphism in social behavior plasticity. Indeed, female flies from select DGRP lines showed baseline SD phenotypes consistent with their male counterparts, yet mechanical injury did not significantly affect female SD (Fig. S4B; also see Fig. 5G). Considering that DSK-expressing MP1 neurons originate from the larval brain (43), we hypothesized that early-life experience is developmentally encoded in DSK neuron activity via the male-specific neural pathway and adaptively expressed for social plasticity in adults.

**Fig. 5.**
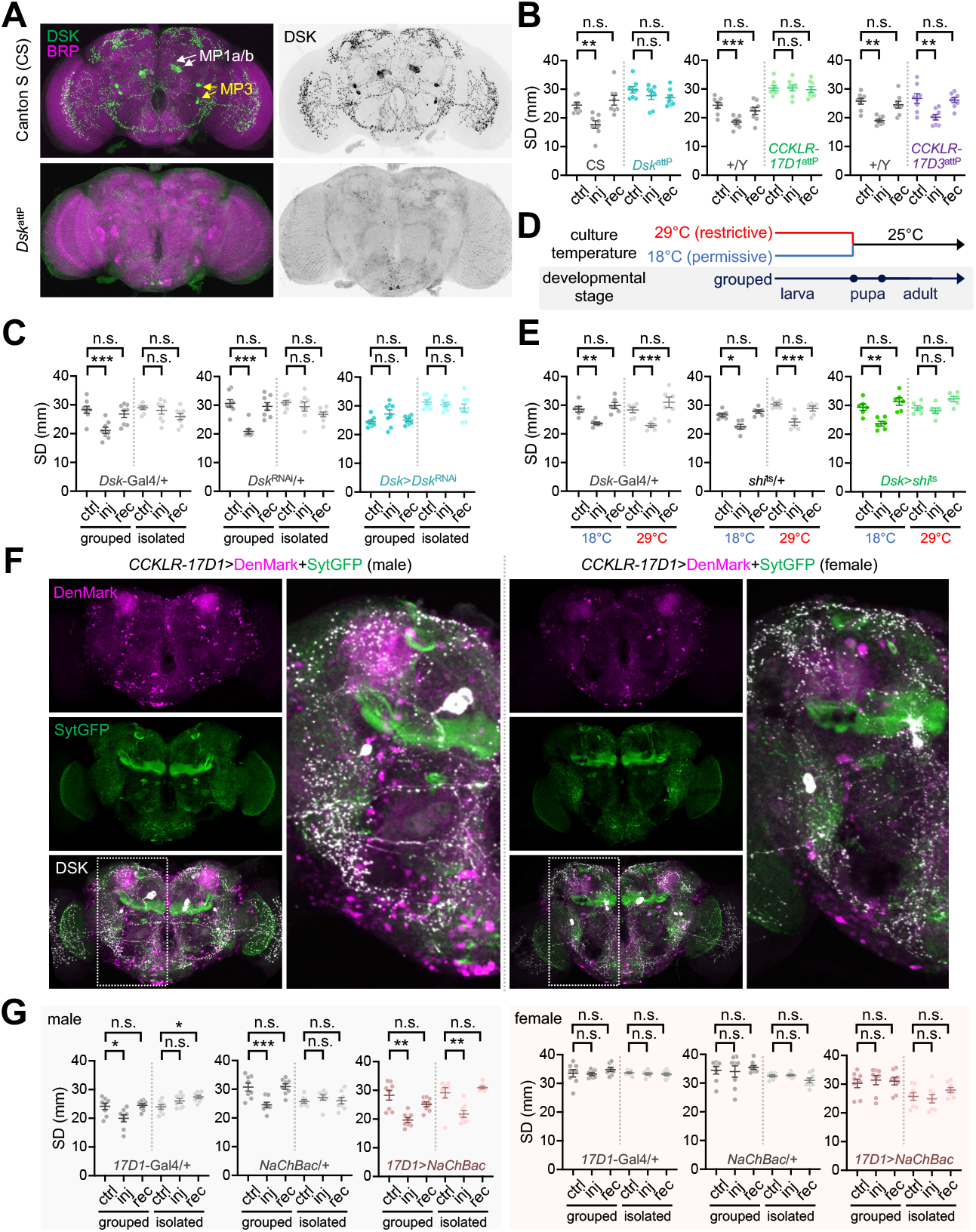
Genetic manipulations of DSK signaling imitate social experience. (A) *Dsk*^attP^ mutant brain expresses barely detectable DSK peptides. (B and C) Genetic silencing of DSK-CCKLR-17D1 signaling by genomic deletions (*Dsk*^attP^ or *CCKLR-17D1*^attP^) or DSK depletion (*Dsk*>*Dsk*^RNAi^) impairs the injury-induced plasticity of SNB. Data represent means ± SEM (*n* = 8). n.s., not significant; ^**^*P* < 0.01, ^***^*P* < 0.001, as determined by 2-way ANOVA with Tukey’s multiple comparisons test. (D and E) Conditional blockade of DSK neuron transmission at larval stage is sufficient to blunt social experience-dependent plasticity of SNB. Transgenic crosses for DSK-specific expression of the temperature sensitive *shibire* allele (*Dsk*>*shi*^ts^) were kept at either restrictive (29°C) or permissive temperature (18°C) for synaptic transmission until the end of larval stage. Data represent means ± SEM (*n* = 6). n.s., not significant; ^*^*P* < 0.05, ^**^*P* < 0.01, ^***^*P* < 0.001, as determined by 2-way ANOVA with Tukey’s multiple comparisons test. (F) CCKLR-17D1 neurons display sexually dimorphic dendrites around MP1 projections. Transgenic male or female brains (*CCKLR-17D1*>DenMark+SytGFP) were immunostained with anti-DSK antibody (white) to visualize dendrites (DenMark, magenta) and axons (SytGFP, green) of neurons expressing CCKLR-17D1-Gal4 knock-in along with DSK neuron projections. (G) Transgenic excitation of CCKLR-17D1 neurons confers injury-induced plasticity of SNB in socially isolated males but not females. Data represent means ± SEM (*n* = 8). n.s., not significant; ^*^*P* < 0.05, ^**^*P* < 0.01, ^***^*P* < 0.001, as determined by 2-way ANOVA with Tukey’s multiple comparisons test.

### Male-specific DSK-CCKLR-17D1 signaling mediates SNB plasticity

To determine whether DSK signaling actually contributes to social behavior plasticity, we examined its loss-of-function effects. Genomic deletion of the *Dsk* locus (*Dsk*^attP^) or DSK depletion by RNA interference (*Dsk*>*Dsk*^RNAi^) abolished injury-induced clustering behaviors (Fig. 5A-5C; Fig. S5). Furthermore, genomic deletion of the *Dsk* receptor *CCKLR-17D1* (*CCKLR-17D1*^attP^) but not *CCKLR-17D3* (*CCKLR-17D3*^attP^) comparably masked injury-induced social interactions (Fig. 5B). We employed a temperature-sensitive allele of *Drosophila* dynamin (*shibire*^ts^) (44) to block synaptic transmission specifically at restrictive temperature (Fig. 5D). The conditional manipulation of larval DSK neurons was sufficient to suppress injury-induced clustering in group-cultured adults (Fig. 5E). The genetic effects of DSK signaling were not consistent on baseline SD but specific to injury-induced social plasticity, implicating the DSK-CCKLR-17D1 signaling in the latter process.

CCKLR-17D1-expressing neurons displayed a gender-specific distribution in the adult brain when their dendrites and axons were visualized using specific transgenic markers (i.e., DenMark and SytGFP, respectively) (Fig. 5F). In particular, male brains expressing the CCKLR-17D1 knock-in transgene showed more signals for cell bodies and dendrites around MP1a axon projections than females (34). This was consistent with the high levels of axonal DSK expression and trans-Tango signals in the same region of male brains. We hypothesized that transgenic activation of DSK-CCKLR-17D1 signaling genetically mimics social experience in developmentally isolated flies. Supporting this idea, transgenic excitation of CCKLR-17D1 neurons was sufficient to confer injury-induced social interactions to isolated male flies (Fig. 5G). By contrast, the same transgenic manipulation of CCKLR-17D1 neurons did not induce social plasticity in females, irrespective of their early-life experience. These findings convincingly provide a neuroanatomical basis for male-specific social plasticity.

## Discussion

Our study demonstrated that conspecific individuals show a wide range of preferences for social distancing, and it has likely co-evolved with their inferior traits as a compensatory mechanism. Physiological challenges (e.g., mechanical injury) may adaptively modify the clustering property of a given group. However, the plasticity of group behaviors requires social experience during development. We propose that a subset of the *Drosophila* brain neurons expressing the neuropeptide DSK serves as a neural substrate for social memory, as supported by the intimate coupling of early-life social experience and SNB plasticity to DSK expression, DSK neuron activity, and post-synaptic signaling.

Distinct social behaviors in *Drosophila* (i.e., aggression and mating) have been commonly mapped to specific pairs of DSK neurons (32, 34, 36). Presynaptic partners of DSK neurons are sexually dimorphic, while their post-synaptic effects are differentially mediated via the two DSK receptor pathways (i.e., CCKLR-17D1 for aggression and SNB plasticity; CCKLR-17D3 for sexual behaviors). Intriguingly, this neural architecture for balancing aggression and mating behaviors is analogously conserved between flies and mammals (45). Furthermore, DSK neuron activity correlates with social dominance (i.e., winner effects from aggression) (34) and group housing (32), but not with mating status (36). Accordingly, the DSK-CCKLR-17D1 pathway meets the necessary criteria for male-specific SNB plasticity.

The phenotypic correlates of aggression and SNB in natural populations (i.e., DGRP lines) are consistent with their common neural locus. However, these findings do not necessarily imply that DSK neurons control the two social behaviors in a similar manner. For instance, DSK excitation promotes aggression in both males and females (34), whereas DSK signaling is unlikely to trigger grouping behaviors per se. What remains to be clarified is how DSK signaling gates SNB plasticity and why this process is missing in female flies. Considering that social hierarchy is established in male fights only (46, 47), we speculate that male-specific DSK pathways include extra circuit modalities for contextual processing of social environments and adaptive social structures. The development of neuroanatomical differences between male and female brains should coincide with the acquisition of gender-specified demands for innate behaviors and physiology during evolution (e.g., a reproductive advantage of male clustering under physiologically challenging conditions).

The mammalian DSK homolog CCK shows sexual dimorphism in brain expression and mating behavior response (48-50). Moreover, CCK activation is implicated in aggression and exploratory behaviors (51, 52). It would thus be interesting to determine whether DSK/CCK signaling indeed represents an ancestral mechanism for social memory, sexually dimorphic social behaviors, and their plasticity.

## Materials and Methods

### Fly stocks

All flies were raised in standard cornmeal-yeast-agar food at 25°C. DGRP lines, Canton-S (BL64349), *rut*^1^ (BL9404), *Dsk*^attP^ (BL84497), *Dsk*^2A-Gal4^ (BL84630), UAS-*Dsk*^RNAi^ (BL25869), *CCKLR-17D1*^attP^ (BL84462), *CCKLR-17D1*^2A-Gal4^ (BL84605), *CCKLR-17D3*^attP^ (BL84463), UAS-myrGFP.QUAS-mtdTomato-3xHA; trans-Tango (BL77124), 20XUAS-IVS-jGCaMP7f (BL80906), and UAS-DenMark, UAS-syt.eGFP (BL33065) were obtained from Bloomington Drosophila Stock Center. UAS-*shi*^ts^ and UAS-NaChBac have been described previously (44, 53).

### SNB analysis

Flies were briefly cold-anesthetized (< 15 s) and then transferred to a circular arena (5.5 cm diameter) filled with 2% agar media. Each arena was video-recorded for 10 min using a cellular phone (Samsung Galaxy Note 8 or Samsung Galaxy Note 20). Time-series coordinates of each fly’s position in the arena were extracted from raw video data using an in-house Python code (https://github.com/jiunbae/tracking-fly). Social distance (SD) was calculated from the position coordinates of individual flies at a given time and averaged over time. The average walking speed of individual flies and the interquartile ranges of the positional centroid in a given group of flies were also calculated over time (https://github.com/KJKwon/2023_FlyBehavior). The centroid velocity was determined from the fourth quarter of video-recording data, given more evident clustering phenotypes at the later period.

### Fly manipulations

*Drosophila* eggs were collected on a plate filled with grape juice media (https://cshprotocols.cshlp.org/content/2007/9/pdb.rec11113). Each egg was gently transferred to an isolation chamber containing 300 ul of cornmeal-yeast-agar food for social isolation. The chamber was sealed using parafilm with a tiny hole for air circulation and kept at 25°C before relevant experiments. For physical injury, the mesothoracic segment of 3-day-old flies was pierced (< 1-mm depth) using a sterilized needle.

### Larval behavior and developmental analyses

For clustering assay, a group of 3rd-instar larvae (n = 20) were loaded onto a cylinder vial [2.3 cm (d) x 9.5 cm (h)] containing standard cornmeal-yeast-agar food. The food vial was divided into 4 sectors. The maximum number of clustering larvae per sector was scored from each vial, and the percentage of clusters with given larvae numbers was calculated from 30 vials. For digging assay, a 2D-arena [2 cm (w) x 0.5 cm (d) x 4 cm (h)] was filled up to 2 cm with standard cornmeal-yeast-agar food. Either a group of 3rd-instar larvae (n = 20) or an isolated 3rd-instar larva was transferred to the digging-assay arena and then allowed to explore it for 12 h before measuring digging depth from the surface (15). For food intake assay, a wider 2D-arena [8 cm (w) x 0.5 cm (d) x 4 cm (h)] was filled up to 2 cm with standard cornmeal-yeast-agar media containing 1% brilliant blue FCF (JUNSEI, 64350-0410). A group of 3rd-instar larvae (n = 20) or an isolated 3rd-instar larva was transferred to the food intake arena and then allowed to explore it for 12 h. Each larva was gently homogenized in 50 ul of distilled water, and the absorbance of individual larval extracts was measured at 627 nm using a microplate reader (Tecan, Infinite M200). Developmental time was measured by the first eclosion day in grouped-egg (n = 50) or isolated-egg cultures (n = 81) per experiment. Male and female progeny numbers were scored from grouped (n = 100) or isolated egg cultures (n = 81) per experiment, and the ratio of males to total flies from each culture was calculated accordingly.

### Maze assay

Yeast paste was placed at the corner of a 14 cm x 14 cm transparent maze, and the maze was put on a white-light box to avoid any phototatic effects. A group of 4 flies was starved for 6 h and then transferred to the maze by an aspirator. A training session was completed when 3 or more pioneer flies reached the food, and the trained flies were transferred to an empty vial containing water only. The pioneer training was repeated three times. For the maze test, a group of 16 flies (16 naive or 12 naive + 4 pioneer flies) was briefly cold-anesthetized (< 15 s) and then placed at the opposite corner of the maze to the yeast paste. The 75% arrival time was recorded when 12 or more flies reached the food.

### Transcriptome analysis

Total RNAs were extracted from fly heads and purified using the PureLink RNA mini kit according to the manufacturer’s instructions (Invitrogen). RNA quality was assessed by Bioanalyzer using the Agilent RNA 6000 pico kit (Agilent Technologies). mRNAs were further enriched using the NEBNext Poly(A) mRNA Magnetic Isolation Module (New England Biolabs). RNA-seq libraries were constructed using the NEBNext Ultra Directional RNA Library Prep Kit for Illumina (New England Biolabs) and subsequently sequenced by Illumina NovaSeq 6000 or Illumina NextSeq 500/550 (LabGenomics, Republic of Korea). RNA-seq reads were processed using trimmomatic (54) (version 0.39) with the default option to remove bases with low-quality scores or from sequencing adapters. The trimmed reads were then mapped to the Drosophila melanogaster reference genome at Flybase (R6.46) using STAR (55) (version 2.7.10b). Differentially expressed genes were determined using DEseq2 (56) (version 1.40.0; >2 fold change with adjusted *P* < 0.05). Transcripts undetectable in more than half of the RNA-seq libraries were excluded from the analysis. Overrepresented gene ontology terms were identified by Fisher’s exact test (false discovery rate < 0.05) with PANTHER (57) (version 17.0).

### Quantitative brain imaging

Whole-mount brain imaging was performed as described previously (58). The primary antibodies used in immunostaining included rabbit anti-DSK (Boster Bio, DZ41371; diluted at 0.5 ug/ml) and mouse anti-BRP antibodies (Developmental Studies Hybridoma Bank, nc82; diluted at 1:1,000). The GCaMP signals from live brains were recorded at room temperature using an FV1000 microscope (Olympus). Fluorescence intensities from regions of interest in confocal images were quantified by background normalization [(S-B)/B] using ImageJ software.

### Statistical analysis

Statistical analyses were performed using GraphPad Prism or R (version 4.2.3). Shapiro-Wilk test was followed by F-test (two samples) or Brown-Forsythe test (multiple samples) to check normality (*P* < 0.05) and equality of variances (*P* < 0.05), respectively. For two-sample comparisons, parametric datasets with equal variance were analyzed by unpaired t-test. For multiple-sample comparisons, 1) parametric datasets with equal variance were analyzed by ordinary ANOVA with Tukey’s multiple comparisons test; 2) parametric datasets with unequal variance were analyzed by Welch’s ANOVA with Dunnett’s T3 multiple comparisons test (1-way) or by aligned ranks transformation ANOVA with Wilcoxon rank sum test (2-way); and 3) nonparametric datasets with equal variance were analyzed by Kruskal-Wallis test with Dunn’s multiple comparisons test (1-way) or by aligned ranks transformation ANOVA with Wilcoxon rank sum test (2-way). For comparisons between repeatedly measured samples, datasets were analyzed by paired t-test (two samples) or 2-way repeated measures ANOVA with Sidak’s multiple comparisons test (multiple samples). Sample sizes and *P* values obtained from individual statistical analyses were indicated in the figure legends.

## Data, materials, and software availability

The datasets generated and analyzed during the current study are available in the European Nucleotide Archive repository (accession number PRJEB61423). The python scripts that support the findings of this study are available from the author’s GitHub webpage under the links https://github.com/jiunbae/tracking-fly and https://github.com/KJKwon/2023_FlyBehavior.

## Acknowledgments

We thank Bloomington Drosophila Stock Center, Developmental Studies Hybridoma Bank, and Korea Drosophila Resource Center for reagents; Kenneth Wilson and Pankaj Kapahi for raw data from their phenotypic DGRP screens. This work was supported by grants from the Suh Kyungbae Foundation (SUHF-17020101[C.L.]); from the National Research Foundation funded by the Ministry of Science and Information & Communication Technology (MSIT), Republic of Korea (NRF-2021M3A9G8022960 [C.L.]; NRF-2018R1A5A1024261 [C.L.]; NRF-2023R1A2C100627511 [T.K.]); from Basic Science Research Program through the National Research Foundation funded by Ministry of Education (NRF-2018R1A6A1A03025810 [T.K.]).

## Author Contributions

Conceptualization, J.J., K.K., T.K., and C.L.; Methodology, J.J. and K.K.; Validation, J.J. and K.K.; Formal Analysis, J.J., K.K., T.K., and C.L.; Investigation, J.J., K.K., T.K.G., and J.L.; Writing - Original Draft, J.J., K.K., T.K., and C.L.; Writing - Review & Editing, T.K. and C.L.; Visualization, J.J., K.K., T.K., and C.L.; Supervision, T.K. and C.L.; Funding Acquisition, T.K. and C.L.

## Figure Legends

**Data S1. Normalized gene expression in individual DGRP lines under grouped vs. isolated culture conditions**.

**Data S2. DEG analyses between distinct social groups (short vs. long SD; or grouped vs. isolated)**.

**Data S3. DEG analyses among individual DGRP lines**.

