## Supplementary material for "*Drosulfakinin* signaling encodes early-life memory for adaptive social plasticity": Fig S1-S5

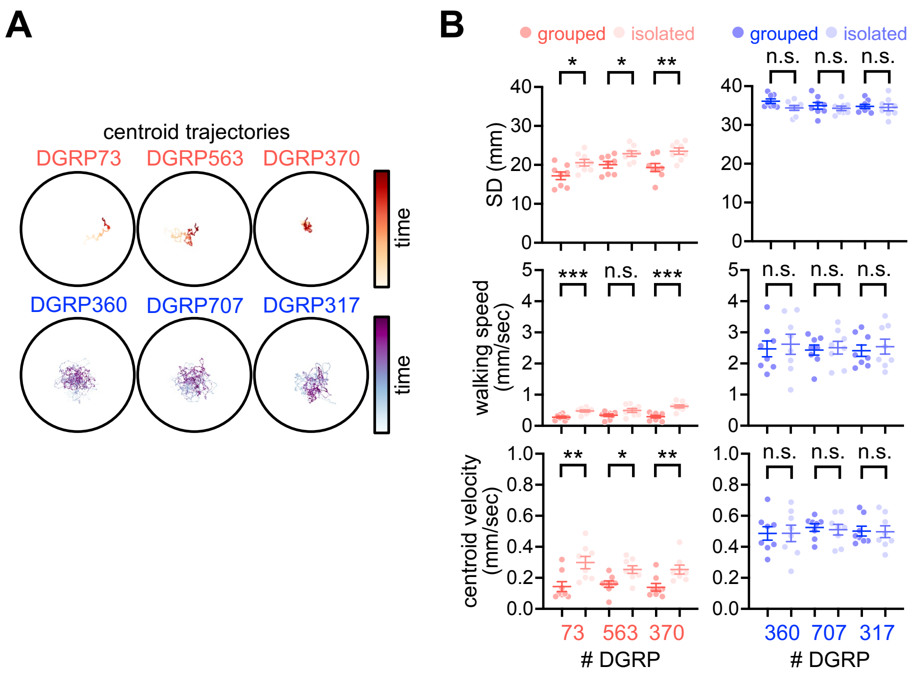


**Fig. S1. Social network behaviors in the three representative DGRP lines displaying short or long SD.** (A) 10-min centroid trajectories of short- (red; DGRP73, 563, 370) or long-SD lines (blue; DGRP360, 707, 317). (B) The short-SD lines exhibit slow walking speeds and low centroid velocities compared to the long-SD lines. Social isolation modestly impaired SNB only in the short-SD lines. Data represent means ± SEM ($n$ = 8). n.s., not significant; *$P < 0.05$, **$P < 0.01$, ***$P < 0.001$, as determined by unpaired t-test.

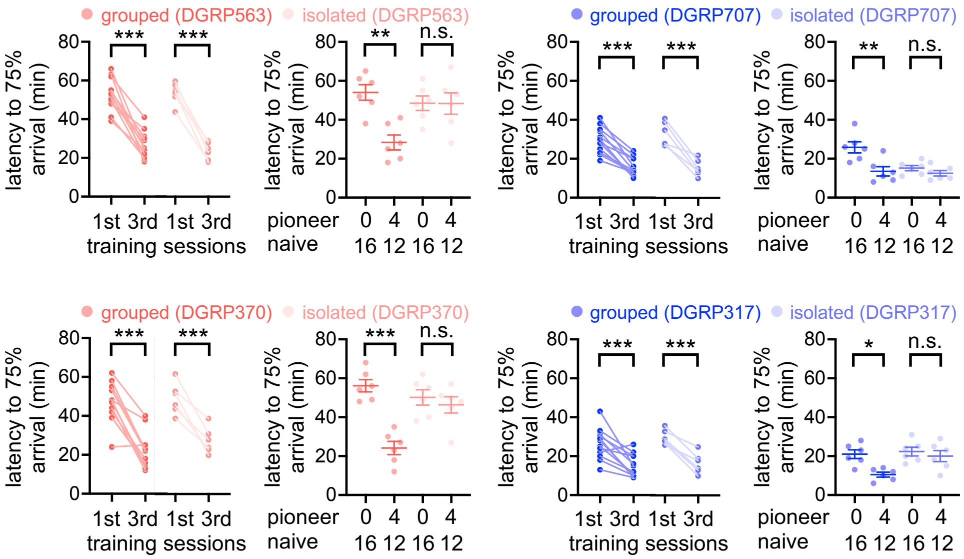


**Fig. S2. Social isolation blunts pioneer effects on food-seeking behaviors in a group of naive flies.** Pioneer groups of short- (red; DGRP563, 370) and long-SD lines (blue; DGRP707, 317) were effectively trained in the maze assay but they failed to improve the efficiency of food-seeking behaviors with groups of socially isolated flies. Data represent means ± SEM ($n$ = 6-12). n.s., not significant; *$P < 0.05$, **$P < 0.01$, ***$P < 0.001$, as determined by 2-way repeated measures ANOVA with Sidak's multiple comparisons test (training sessions) or 2-way ANOVA with Tukey's multiple comparisons test (test sessions).

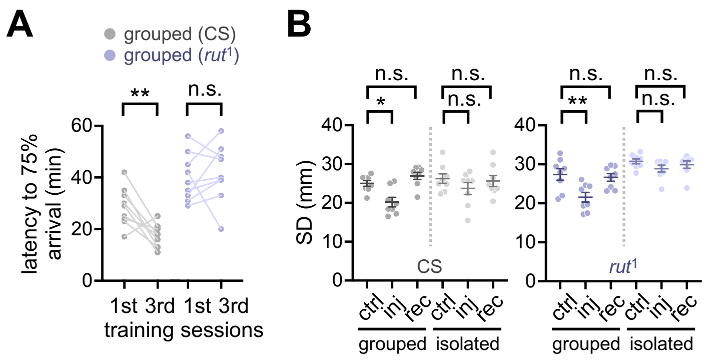


**Fig. S3. *rutabaga*-dependent working memory is dispensable for social memory.** (A) The plasticity mutant *rutabaga* (*rut*$^1$) fails to improve food-seeking behaviors in the maze after repetitive trainings. Data represent means ± SEM (*n* = 9). n.s. not significant; ***P* < 0.01, as determined by paired t-test. (B) *rut* mutants display injury-induced social plasticity when socially enriched but not socially isolated. Data represent means ± SEM (*n* = 8). n.s., not significant; **P* < 0.05, ***P* < 0.01, as determined by 2-way ANOVA with Tukey's multiple comparisons test. CS, Canton-S (wild-type) flies.

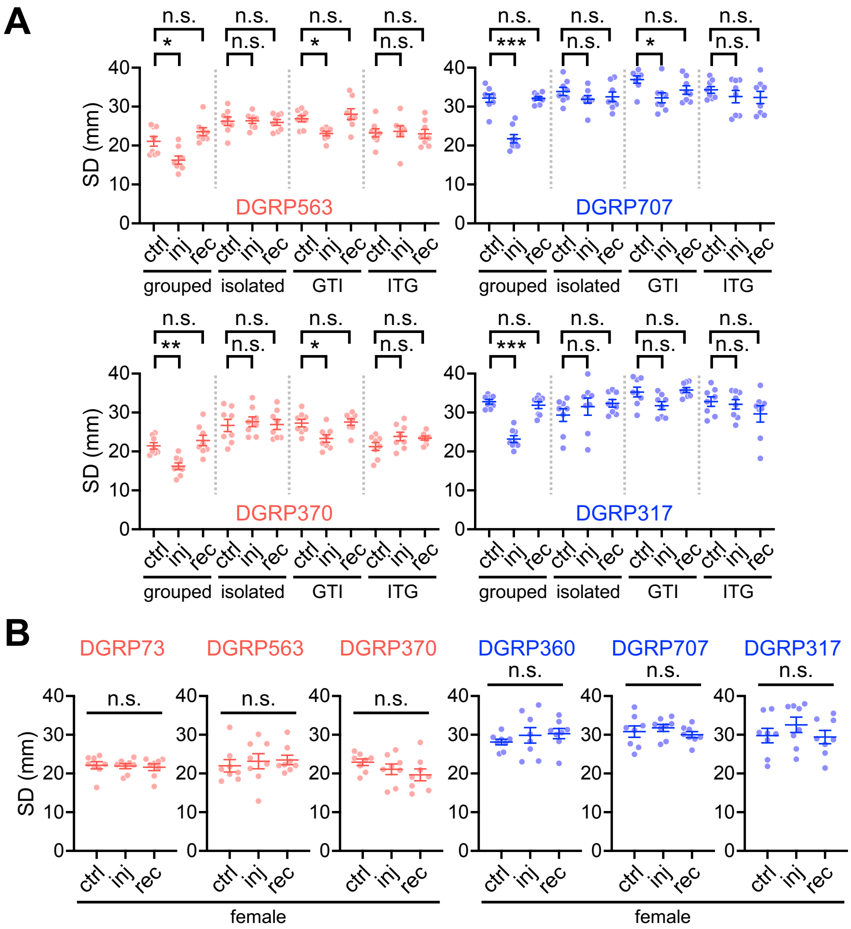


**Fig. S4. Early-life social experience is necessary for social behavior plasticity in male flies.** (A) Physical injury induced clustering behaviors in group cultured but not developmentally isolated flies. Data represent means ± SEM ($n$ = 8). n.s., not significant; *$P$ < 0.05, **$P$ < 0.01, ***$P$ < 0.001, as determined by 1-way ANOVA with Tukey's multiple comparisons test. GTI, grouped-to-isolated culture transition; ITG, isolated-to-grouped culture transition. (B) Female flies did not display injury-induced social plasticity. SD was measured in group-cultured female flies from individual DGRP lines. Data represent means ± SEM ($n$ = 8). n.s., not significant as determined by 1-way ANOVA with Tukey's multiple comparisons test.

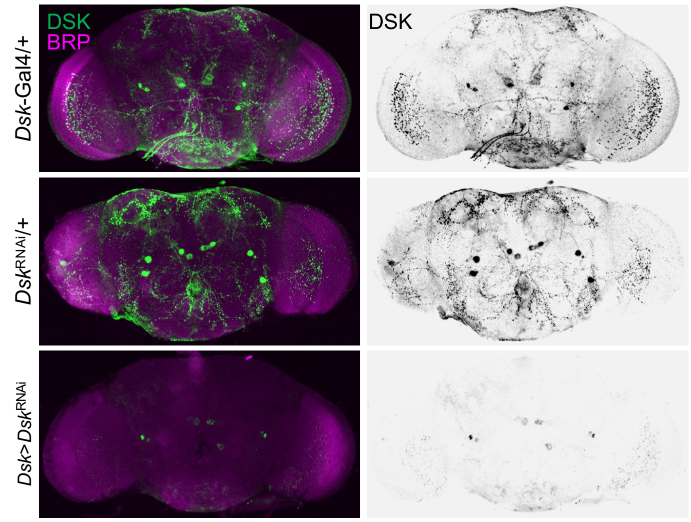


**Fig. S5. Transgenic *Dsk* RNAi effectively depletes DSK peptides in the adult brain.** The *Dsk* RNAi transgene was expressed in DSK neurons by the *Dsk* knock-in Gal4 driver (*Dsk*>*Dsk*$^{RNAi}$). Whole-mount transgenic brains were co-immunostained with anti-DSK (green) and anti-BRP (magenta) antibodies. Heterozygous flies for Gal4 (*Dsk*-Gal4/+) or RNAi transgenes (*Dsk*$^{RNAi}$/+) served as negative controls.
